# LapTrack: Linear assignment particle tracking with tunable metrics

**DOI:** 10.1101/2022.10.05.511038

**Authors:** Yohsuke T. Fukai, Kyogo Kawaguchi

**Affiliations:** Nonequilibrium Physics of Living Matter RIKEN Hakubi Research Team, RIKEN Center for Biosystems Dynamics Research, 2-2-3 Minatojima-minamimachi, Kobe, 650-0047, Japan; RIKEN Cluster for Pioneering Research, 2-2-3 Minatojima-minamimachi, Kobe, 650-0047, Japan; Universal Biology Institute, The University of Tokyo, 7-3-1 Hongo, Bunkyo-ku, Tokyo, 113-0033, Japan

## Abstract

**Motivation:** Particle tracking is an important step of analysis in a variety of scientific fields, and is particularly indispensable for the construction of cellular lineages from live images. Although various supervised machine learning methods have been developed for cell tracking, the diversity of the data still necessitates heuristic methods that require parameter estimations from small amounts of data. For this, solving tracking as a linear assignment problem (LAP) has been widely applied and demonstrated to be efficient. However, there has been no implementation that allows custom connection costs, parallel parameter tuning with ground truth annotations, and the functionality to preserve ground truth connections, limiting the application to datasets with partial annotations.

**Results:** We developed LapTrack, a LAP-based tracker which allows including arbitrary cost functions and inputs, parallel parameter tuning, and ground-truth track preservation. Analysis of real and artificial datasets demonstrates the advantage of custom metric functions for tracking score improvement. The tracker can be easily combined with other Python-based tools for particle detection, segmentation, and visualization.

**Availability and implementation:** LapTrack is available as a Python package on PyPi, and the notebook examples are shared at https://github.com/yfukai/laptrack. The data and code for this publication are hosted at https://github.com/NoneqPhysLivingMatterLab/laptrack-optimization.

## I. INTRODUCTION

Automated tracking of particles in timelapse images is important in a wide range of fields in science, and is especially crucial in creating large datasets of cell lineages in biological studies. Recently there has been considerable development in tracking algorithms, where methods based on supervised machine learning are increasingly being developed [1–3]. The diverse nature of live imaging tasks, however, frequently requires tracking without large-scale ground-truth annotations, emphasizing the need for a robust tracking algorithm with a small number of parameters that can be tuned by manual annotations.

Defining and optimizing a global cost function to appropriately penalize wrong connections is a common approach in robust tracking methods. If the cost function is a linear sum of the costs associated to connections, we can employ efficient algorithms [4] to solve the global optimization problem called the linear assignment problem (LAP). The LAP-based tracking method has proven to be accurate and robust, especially for data with higher particle density. To deal with particle splitting (by division or over-segmentation) or merging (by under-segmentation), which is common in the data of live-imaged cells, [5] further developed a two-stage LAP method, with the second stage dedicated to the connection of splitting and merging branches. The cost function in their case was the squared Euclidean distance between the positions of the objects, with additional intensity-associated costs for splitting and merging.

Tools have been developed to provide similar LAP-based algorithms with splitting and merging detection; TrackMate [6, 7], for example, provides distance-based LAP-based tracking with particle detection and segmentation workflow and a method to conduct manual correction, all within the Java-based framework in ImageJ [8, 9]. Cell-ACDC [10], which was originally designed for yeast analysis, also implements an overlap-based LAP tracker with splitting detection, as well as various functions ranging from image alignment to manual correction that support the entire analysis workflow in Python.

Although other highly accurate methods have been proposed to work for the tracking problem with cell divisions [11], no single tracking algorithm is likely to be perfect for the diverse experimental situations. To obtain near-perfect segmentation and tracking for specific data, users must still optimize the segmentation and tracking steps, automatically or manually. In this regard, the LAP-based algorithm that robustly works with a small number of parameters continues to play a key role in generating the initial tracking data without large-scale manual annotation.

An adaptive improvement to the original LAP-based tracking with distance can be made by using additional features taken from the cell images. For example, we can extract the morphology of each cell such as its shape and size from typical live cell images, as well as the signal levels from multiple fluorescent channels. The consistency of cell shape and fluorescent signals across time frames is useful when tracking is conducted by human eyes, especially when the frame rate of the data is not high enough. Therefore, it is desirable to be able to implement arbitrary inputs and cost functions in the LAP-based tracking scheme, as well as to tune the parameters using partial ground-truth annotations.

These requirements motivated us to build a tool that recapitulates the LAP algorithm [5, 6] with additional flexibility and modularity; LapTrack is designed as a simple intermediate in the whole tracking pipeline that takes positions and features of the particles and returns the LAP-optimized tracks. The three unique features of LapTrack are: (1) arbitrary tunable cost functions for particle connection, (2) integratability with other Python tools, and (3) the functionality to preserve the ground-truth (annotated) connections. Within this framework, we can implement user-defined cost functions for connections that can take an arbitrary number of inputs. The tracking function is modularized and documented as an API so that it can be integrated into any custom workflow in Python, allowing parallel parameter optimization as well as visualization of results in easy steps.

In this paper, we demonstrate how this pipeline can be used not only to optimize the tracking in a supervised manner, but how it is also useful for efficient manual correction of the tracks when combined with visualization tools such as napari [12].

## II. METHODS

### A. Datasets

We here describe the data that we used to demonstrate the use cases of LapTrack: live cell images with ground truth segmentation and tracking (mouse paw epidermis and cell migration) as well as simulated data (coloured particles), provided in https://github.com/yfukai/laptrack-optimization. We also used the high-density vesicles, yeast, and 3D *Drosophilla* data to show that the tracking pipeline works for a broad range of applications.

#### 1. Mouse paw epidermis

The segmentation data and ground truth tracking result collected and analyzed in [13, 14] were used for benchmark. The dataset contains 236 to 327 cells in the observation area.

#### 2. Cell migration

The images, segmentation data for a portion of frames, and the ground truth tracking result were downloaded from Zenodo [15]. Segmentation was conducted by Cellpose [16] and manually corrected in napari [12]. The ground truth tracking result was also manually validated and corrected. The dataset contains 218 to 434 cells in the 648.95 μm × 648.95 μm observation area.

#### 3. Coloured particles

We simulated the Brownian motion of four-hundred particles with colours in a two-dimensional box of size 20 × 20 with periodic boundary conditions. The particles were split into two species, *a* and *b,* where the interaction between the particles was set as harmonic repulsion with the spring constants set as 1 for a and a pairs, 1.2 for a and b pairs, and 1.4 for b and b pairs. The dynamics was simulated with the `simulate.brownian` routine in Jax-MD [17] with the parameters *kT* = 0.1 and *dt* = 0.001, where the mass and friction coefficient were set to the default values, 1 and 0.1. For each particle, labeled by *i,* a random integer *n_i_* between 0 to 7 is assigned. The *feature vector c_i_* ∈ ℝ^3^, corresponding to RGB colours, of each particle at each time step is then assigned as 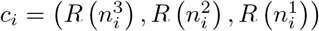, where 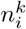 is the *k*-th digit of *n_i_* in the binary representation and 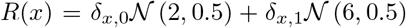, where 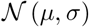 is the normal random variable with mean *μ* and the standard deviation *σ*. When used for the tracking benchmark, the particles crossing the boundary are regarded as disconnected and belong to different tracks.

#### 4. Demonstration

The simulated single-molecule dataset was downloaded from the Particle Tracking Challenge website http://bioimageanalysis.org/track/ [18]. We used the high-density vesicles data set with *SNR* = 7. The blobs were detected by the Laplacian-of-Gaussian detector, `skimage.feature.blob_log` function in scikit-image [19], with the parameters `min_sigma=1,max_sigma=5, num_sigma=5 and threshold=0.05`. The detected points were tracked by `LapTrack` with `track_cost_cutoff=100`.

The yeast dataset was downloaded from the Yeast Image Toolkit website http://yeast-image-toolkit.org/. The data in IT-Benchmark2/TestSet4/RawData were segmented by Cellpose 0.7.2 [16] with the parameters `model_type=“cyto”`, `net_avg=True`, and `diameter=30` in the eval function. The centroids of each segmented region were tracked by `LapTrack` with the default metric and `track_cost_cutoff=100, splitting_cost_cutoff=2500`.

The 3D *Drosophilla* dataset (Fluo-N3DH-CE) was downloaded from the Cell Tracking Challenge website http://celltrackingchallenge.net/ [11]. The data included marked cell positions in each time frame, which were connected to generate tracks by `LapTrack` with `track_cost_cutoff=10000,splitting_cost_cutoff=2500`.

### B. Tracking implementation

The implemented particle tracking algorithm follows the method proposed in [5], with modifications following TrackMate [6, 7] and additional flexibility as we describe in the following sections.

#### 1. Frame-to-frame LAP

In the first step, the points in successive frames are connected by solving LAP, and then generating tracks without splits and merges [Fig. 1(a) left top]. Specifically, for every pair of points with properties (such as Euclidean coordinates) *x_i_* and *x_j_* at frames *t* and *t*+1, the costs *l_ij_* = *l*(*x_i_, x_j_*) are computed by a user-definable metric function *l.* Costs *d* and *b* are then assigned to the particles not connected to any of the particles in the next and previous timesteps, respectively. The optimal assignment is found by minimizing the cost [5]:

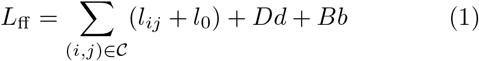

where 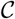 is the set of all connected index pairs, *B* and *D* are the number of the points which does not have the connection to the previous and next timesteps, respectively, and *l*_0_ = min (*l_ij_, d,b*) (See Supplementary Material for algorithm details). In the default setting, *d* and *b* are calculated as 1.05 × *c*^90%^, where *c*^90%^ is the 90% percentile value of the all finite entries in {*l_ij_*}_*ij*_ [5]. The default metric for *l* is set to the squared Euclidean distance 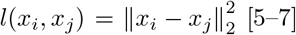 [5–7] with which the cost-minimizing association can be interpreted as the maximum log-likelihood solution for Brownian particles when we ignore splitting and merging [20].

**FIG. 1.**
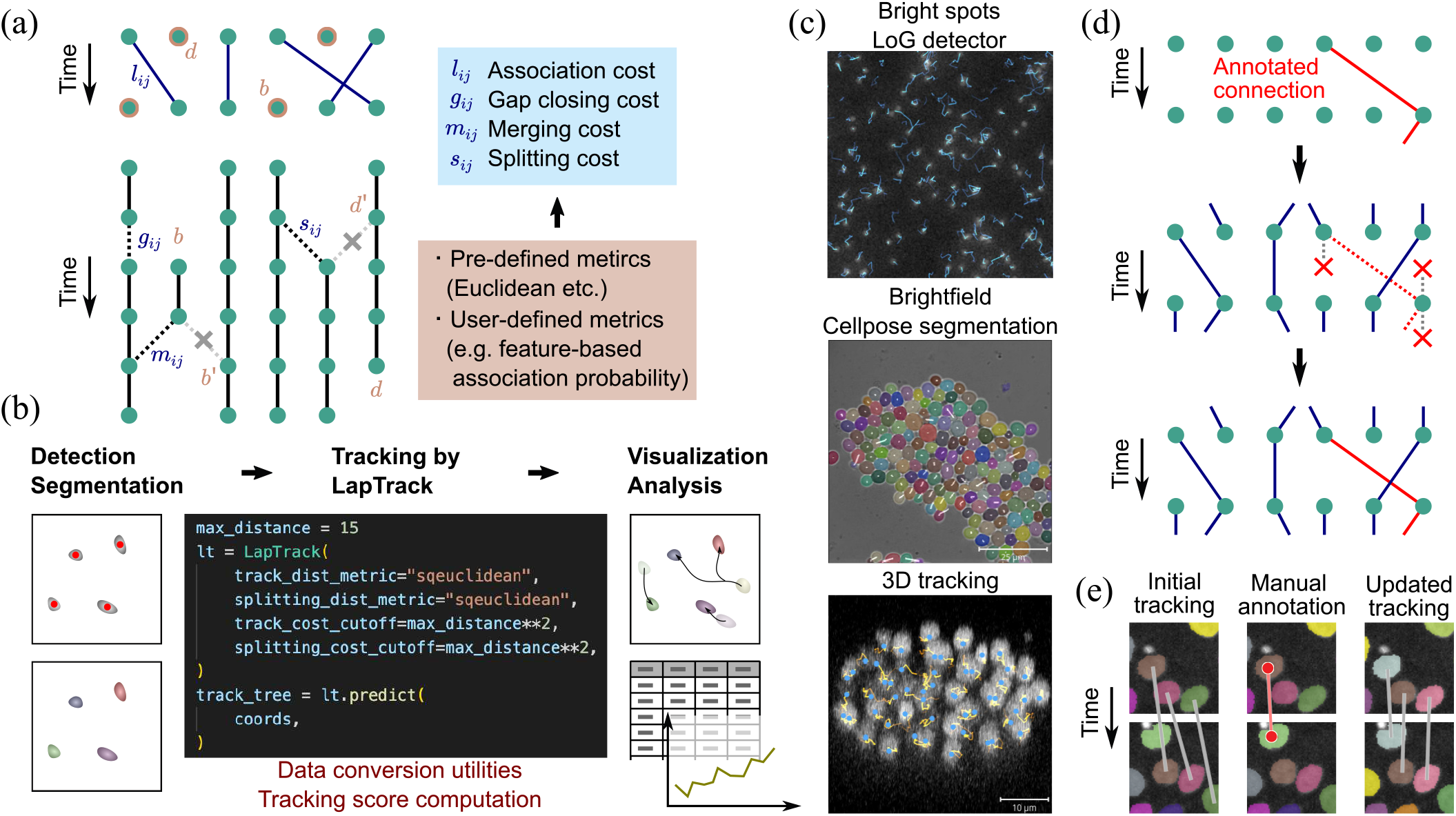
(a) The schematic for the tracking algorithm (See main text). (b) Expected workflow for cell segmentation, tracking and analysis using tools in Python. The particle detection or segmentation results can be directly supplied to LapTrack. The tracking result can be directly visualized and analyzed in Python. (c) Examples of tracks generated by LapTrack. The lines indicate the result tracks. (top) The dataset from Particle Tracking Challenge, detected by the Laplacian of Gaussian detector. (middle) The dataset from the Yeast Image Toolkit website, detected by Cellpose. (bottom) The C.elegans developing embryo dataset from the Cell Tracking Challenge website. (d) The schematic for the tracking algorithm with freezing annotated connections. (top) Annotated connections (red lines). (middle) Connections from (to) a point that has an annotated connection from (to) itself are forbidden. (bottom) The verified connections are added to the tracking tree. The split and merges are treated similarly. (e) Illustration of the manual-correction-aware tracking with napari (See main text) using the cell migration dataset. (left) Original tracking result with mistakes (gray lines). (middle) Annotation points are added in napari (red points) to specify a correct connection (red line). (right) Updated tracking result after annotation. The annotated track as well as tracks nearby are automatically corrected (gray lines).

#### 2. Segment-connecting LAP

In the second step, another LAP is solved to predict splitting, merging, and gap closing [Fig. 1(a) left bottom]. Gap closing connects free segment ends with allowing frame skips. The gap closing cost *g_αβ_* = *g*(*x_α_, x_β_*) is calculated by a user-definable metric *g* for all possible connections between free ends up to a specified frame difference, and the splitting and merging costs *s_α β_* = *s*(*x_α_, x_β_*) and *m_αβ_* = *m*(*x_α_, x_β_*) are calculated for all possible connections between a free end and a track midpoint by user-definable metrics *s* and *m.* The metrics *g, s,* and *m* default to the squared Euclidean distance. Then the optimal assignment is calculated by minimizing the overall cost

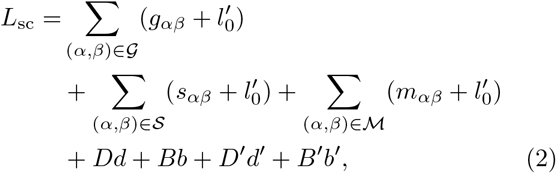

where 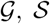 and 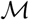 are the set of all gap-closing, splitting and merging index pairs, *D* and *B* are the number of the unconnected track ends and starts, *D*′ and *B*′ are the number of the track middle points that are not connected to other track ends as the split or merge (costs *d*′ and *b*′ are assigned to them, respectively), and 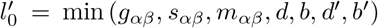 (See Supplementary Material for details). In the default setting, *d*, *b*, *d*′, and *b*′ are calculated analogously to the frame-to-frame LAP.

#### 3. Freezing annotated tracks

We implemented an option to specify partial tracks within the data to be fixed as ground-truth verified connections [Fig. 1(d)]. Fixing the correct tracks is especially useful when conducting manual corrections using visualization tools such as napari. As we demonstrate[21] [Fig. 1(e)], track connections can be specified to be fixed by annotating the cell regions before re-running the LAP-based tracking. The resulting track preserves the training data tracks due to the masking scheme [Fig. 1(d)].

#### 4. Parameter optimization

In practice, we introduce the cutoff for the costs, *g_αβ_*, *s_αβ_* and *m_αβ_*, above which those values are regarded as infinity. The values of the cutoffs can affect the performance as demonstrated in Sec. III A, but it is difficult to optimize those values due to the non-differentiablity of the LAP algorithm [22] and the high computational cost for repeating the tracking routine. We therefore used non-gradient optimization methods to optimize the specified sets of the parameters in parallel using the package Ray Tune [23] with the Optuna optimizer [24] and random search. We selected the parameters that achieved the highest connection Jaccard index value or true positive rate, depending on the type of the training data (Sec. IIC1).

#### 5. Analysis pipeline

LapTrack is written in Python with explicit API documentation and can be integrated with, for example, particle detectors in scikit-image and deep-learning-based segmentation packages such as Cellpose [16] [Fig. 1(b,c)]. The output data is the networkx [25] directed tree, which can be analyzed using network analysis functions in the package. We also implemented utilities to convert data into pandas dataframes [26, 27]. In this paper, we used the ground truth segmentation for each dataset as the input, and analyzed the result tracks by networkx and pandas. The tracking and analysis Python scripts are provided at https://github.com/NoneqPhysLivingMatterLab/laptrack-optimization.

### C. Metrics for the tracking results

To measure the performance of tracking, we employed the following metrics, which can also be calculated within LapTrack.

#### 1. Overall tracking scores

To measure the overall track consistency, we calculated the *target effectiveness* (TE) and *track purity* (TP) [1, 28], which penalize the false negative and the false positive detections, respectively. Let us denote the set of the ground truth tracks by 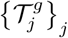. and the predicted tracks by 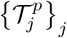. The TE for a single ground truth track 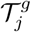 is calculated by finding the predicted track 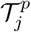 that overlaps with 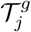 in the largest number of the frames, and then dividing the overlap frame counts by the total frame counts for 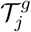. The TE for the total dataset is calculated as the mean of TEs for all ground truth tracks, weighted by the length of the tracks. The track purity is analogously defined with 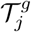 and 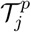 being swapped in the definition. We also measured the *mitotic branching correctness* [1, 28], defined as the fraction of the number of correctly detected divisions over the total number of the divisions.

#### 2. Overlap between predicted and ground truth connections

During the parameter optimization, we used a less computationally expensive quantity, the *Jaccard index* and the *true positive rate* of the connections to measure how well the predicted connections overlap with the ground truth. ‘The quantity is defined by 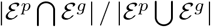 and 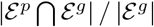, respectively, where we denoted the set of predicted and ground-truth connections by 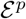 and 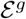, respectively, and the size of a set 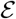 by 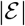.

## III. RESULTS

### A. Distance cutoffs can be optimized to increase performance

We first investigated the performance against varied cost cutoffs in the simplest cases where the costs for the connecting, gap closing, and splitting are the squared Euclidean distance between the centroids. Specifically, we varied the maximum distance allowed for frame-to-frame particle association (`max_distance`) and splitting and gap-closing association (`splitting_max_distance`), which defines the cutoff for *l_ij_* and *s_αβ_* (*g_αβ_*), respectively, and investigated how the overall performance changes. In the mouse epidermis dataset [Fig. 2(a)], we conducted grid-search in the parameters `max_distance` and `splitting_max_distance`. We found that there exists a maxima in the TE around some finite length scale, suggesting that optimization is useful in performance improvement even for the cutoff parameters [Fig. 2(b)]. We also found that the correlation of the tracking scores between mouse epidermis data from different regions are high upon changing of the parameters (*r* = 0.96 (*r* = 0.90) for TE (TP) using data with TE > 0.75 (TP > 0.75), respectively (Supplementary Material Fig. S1)), meaning that the optimized parameters are transferable within similar data.

**FIG. 2.**
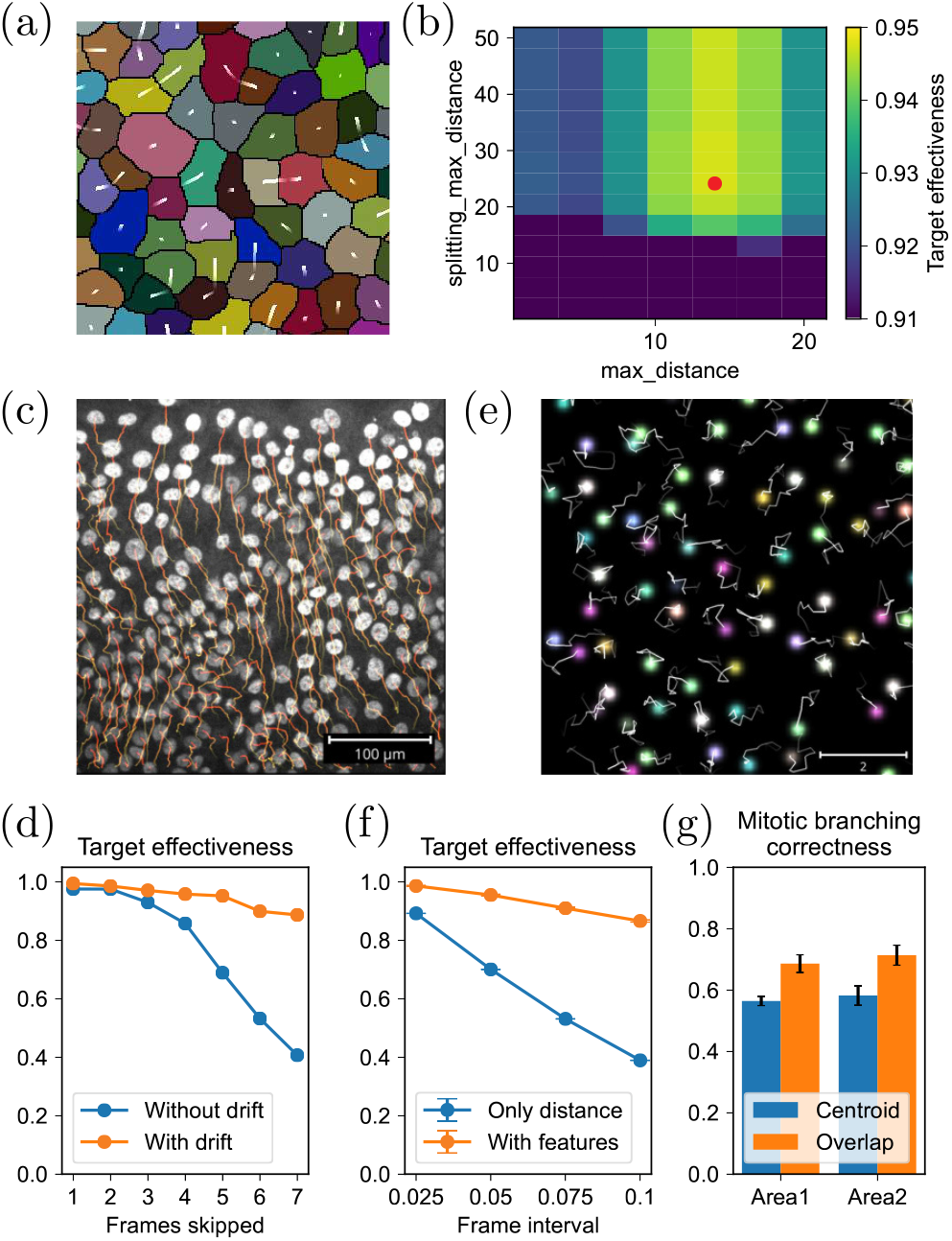
(a) An example snapshot for the mouse epidermis dataset. The white lines indicate the centroid displacement between frames. (b) TE as a function of `max_distance` and `splitting_max_distance` for the mouse epidermis dataset. (c) An example snapshot for the cell migration dataset. (d) TE score for the cell migration dataset with skipped frames, with or without the drift term in the metric. (e) An example snapshot for the coloured particles dataset. (f) TE score for the coloured particles dataset with different frame intervals, with or without the feature difference term in the metric. The error bar indicates the standard deviation of 5 trials. (g) Mitotic branching correctness score for the mouse epidermis dataset, tracked with the centroid distances (centroid) or the overlap ratio (overlap). The error bar indicates the standard deviation of 5 trials.

### B. Tunable cost function improves tracking performance

We next investigated if variable cost functions help improve the tracking score for different datasets.

In Fig. 2(c) we show a snapshot of the cell migration dataset. Here, the cells are moving collectively toward the upper open region. Due to this drift, the LAP-based tracking based solely on Euclidean distance fails especially for large timesteps, as demonstrated in Fig. 2(d) using datasets with skipped frames. This situation can be easily fixed by changing the cost function by adding a drift term to the Euclidean distance as

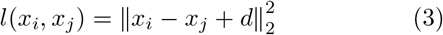

with the drift parameter *d* ∈ ℝ^2^ and defining *g* and *s* analogously [Fig. 2(d), Supplementary Material Fig. S2]. We used 5% of the non-dividing and dividing connections to tune d as well as the cutoffs so that they optimize the true positive rate of the connections. The details are summarized in the Supplementary Material.

Particles may have features to help identify the species, such as sizes and fluorescent intensities. In those cases, we can use those features in addition to the Euclidean distances to improve the performance. To illustrate this, we measured the tracking performance for simulated particles with 8 species, characterized by different sets of feature values corresponding to RGB colours (See II A 3 for details). We then defined the cost function as

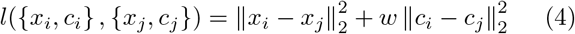

where *c_i_, c_j_* ∈ ℝ^3^ are the feature vectors. We tuned the parameter *w* as well as the distance cutoff using the training data with 100 frames so that the tracking result maximizes the connection Jaccard index. We then measured the tracking scores for an independent dataset with 100 frames. As shown in Fig .2(f), with the features used in the metric, the target effectiveness with large frame interval remains above 0.8 while it drops to ~ 0.4 when only Euclidean distance is in the metric (*w* = 0), illustrating the performance improvement by including the particle features. We also observed improvement of other scores (Supplementary Material Fig. S3).

For segmented images, we can also use the overlap between segmented regions to calculate the cost [7, 10, 29]. The flexible implementation allows us to integrate the overlap metric in addition to the distance in the LAP framework.We defined *l* (with *g* and s analogously) as

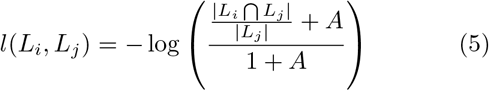

which measures the overlap, where *L_i_* and *L_j_* are the set of pixel coordinates of the segmentation area for the particle *i* and *j* and *A* is a parameter. By comparing the tracking performance for the mouse epidermis dataset with the squared centroid Euclidean distance cases, we found that replacing the metric improves the mitotic branching correctness by ~ 10% [Fig. 2(g)].

## IV. CONCLUSION

In this paper, we showed how the LAP-based tracking pipeline with additional flexibility and optimizability can be useful in improving tracking performance in certain situations, can be easily combined with visualization tools to conduct manual corrections. LapTrack, in large part, is complementary to TrackMate [7], which has a useful GUI and its own optimization pipeline. Compared with TrackMate, LapTrack can take arbitrary inputs and cost functions and is flexible in its output, making it easier to connect with other upstream and downstream analysis pipelines. The tracking function in LapTrack is designed to help making accurate and validated tracks quickly and efficiently, with hope to increase the amount of ground-truth data that can be used in training more sophisticated tracking methods.

With a sufficient amount of manually annotated ground-truth data, machine learning-based approaches will likely outperform the current parameter optimization strategy of simple affinity metrics. Due to its flexibility, our package can be readily combined with strategies such as one-to-one association affinity learning [30, 31], structured learning [2], and the metric learning approach combined with graph neural networks [32], serving as a reusable platform for implementation.

## ACKNOWLEDGEMENTS

We appreciate Dominik Waibel, Benedikt Mairhoermann, Tingying Peng, Carsten Marr and Matthias Meier for discussion, and Rory Cerbus, Somayeh Zeraati, and Takaki Yamamoto for the reading of the manuscript. We acknowledge support by the RIKEN Information systems division for the use of the Supercomputer HOKUSAI Big-Waterfall.

