## Supplementary Text 1 and 2, Supplementary Figures S1-S3 for "LapTrack: Linear assignment particle tracking with tunable metrics"

### SUPPLEMENTARY TEXT 1: LAP IMPLEMENTATION

The tracking algorithm is formulated following [1].

#### Frame-to-frame LAP

Let us consider the points in frame $t$ be $x_i (i = 1 \dots M)$ and the points in frame $t+1$ be $y_i (i = 1 \dots N)$. Let

$$\mathrm{diag}^{\infty}(a)_{ij} := \begin{cases} a & (i = j) \\ \infty & (\text{otherwise}). \end{cases} \tag{S1}$$

We define the cost matrix $C^{\text{ff}}$ by

$$C^{\text{ff}} = \begin{pmatrix} L & \mathrm{diag}^{\infty}(d) \\ \mathrm{diag}^{\infty}(b) & L' \end{pmatrix} \tag{S2}$$

where $L$ and $L'$ are $M \times N$ and $N \times M$ matrices, respectively, defined by

$$L_{ij} = \begin{cases} l_{ij} & (l_{ij} < \hat{l}) \\ \infty & (\text{otherwise}) \end{cases} \tag{S3}$$

and

$$L'_{ji} = \begin{cases} l_0 & (L_{ij} < \infty) \\ \infty & (\text{otherwise}) \end{cases} \tag{S4}$$

where $l_{ij} = l(x_i, y_j)$, $l_0 = \min(L_{ij}, b, d)$, and $\hat{l}$ is the cost cutoff (`track_cost_cutoff`).

We then solve the LAP by the LAPJVsp algorithm [2] implemented as `min_weight_full_bipartite_matching` function in SciPy [3] to minimize the overall cost

$$L_{\text{ff}} = \sum_{i,j} C^{\text{ff}}_{ij} A_{ij} \tag{S5}$$

where $\{A_{ij}\}$ is the assignment matrix satisfying

$$A_{ij} \in \{0, 1\} \tag{S6}$$

$$\sum_i A_{ij} = 1 \tag{S7}$$

$$\sum_j A_{ij} = 1. \tag{S8}$$

The points $x_i, y_j$ are regarded as connected when $A_{ij} = 1$.

#### Segment connecting LAP

Let the track segments generated in the frame-to-frame LAP be $z_\alpha (\alpha = 1 \dots K)$ and the first (last) frame and the coordinates of $z_\alpha$ be $t^{(s)}_\alpha$ $(t^{(e)}_\alpha)$ and $z^{(s)}_\alpha$ $(z^{(e)}_\alpha)$, respectively. Let the frames and the coordinates of the all middle points of the track segments be $t_\alpha$ and $w_\alpha (\alpha = 1 \dots L)$.

Similarly, the cost matrix $C^{\text{sc}}$ is defined by

$$C^{\text{sc}} = \begin{pmatrix} G & M & \mathrm{diag}^{\infty}(d) & \infty \\ S & \infty & \infty & \mathrm{diag}^{\infty}(d') \\ \mathrm{diag}^{\infty}(b) & \infty & \infty & S' \\ \infty & \mathrm{diag}^{\infty}(b') & M' & G' \end{pmatrix} \tag{S9}$$

where the entries are defined in the following.

$G$, $M$, and $S$ are $K \times K$, $K \times L$ and $L \times K$ matrices, respectively, defined by

$$G_{\alpha\beta} = \begin{cases} g_{\alpha\beta} & (g_{\alpha\beta} < \hat{g} \text{ and } t^{(e)}_\alpha < t^{(s)}_\beta \le t^{(e)}_\alpha + \Delta t) \\ \infty & (\text{otherwise}), \end{cases} \tag{S10}$$

$$M_{\alpha\beta} = \begin{cases} m_{\alpha\beta} & (m_{\alpha\beta} < \hat{m} \text{ and } t_\beta = t^{(e)}_\alpha + 1) \\ \infty & (\text{otherwise}), \end{cases} \tag{S11}$$

$$S_{\alpha\beta} = \begin{cases} s_{\alpha\beta} & (s_{\alpha\beta} < \hat{s} \text{ and } t_\alpha = t^{(s)}_\beta - 1) \\ \infty & (\text{otherwise}), \end{cases} \tag{S12}$$

where $g_{\alpha\beta} = g(z^{(e)}_\alpha, z^{(s)}_\beta)$, $m_{\alpha\beta} = m(z^{(e)}_\alpha, w_\beta)$, $s_{\alpha\beta} = s(w_\alpha, z^{(s)}_\beta)$, $\Delta t$ is the maximum allowed gap size (`gap_closing_max_frame_count`), and $\hat{g}$, $\hat{m}$, and $\hat{s}$ are the cost cutoffs (`gap_closing_cost_cutoff`, `merging_cost_cutoff`, and `splitting_cost_cutoff`).

$G'$, $M'$, and $S'$ are defined by

$$G'_{\beta\alpha} = \begin{cases} l'_0 & (G_{\alpha\beta} < \infty) \\ \infty & (\text{otherwise}), \end{cases} \tag{S13}$$

$$M'_{\beta\alpha} = \begin{cases} l'_0 & (M_{\alpha\beta} < \infty) \\ \infty & (\text{otherwise}), \end{cases} \tag{S14}$$

and

$$S'_{\beta\alpha} = \begin{cases} l'_0 & (S_{\alpha\beta} < \infty) \\ \infty & (\text{otherwise}), \end{cases} \tag{S15}$$

where $l'_0 = \min(G_{\alpha\beta}, M_{\alpha\beta}, S_{\alpha\beta}, b, d, b', d')$.

The LAP is solved in the same way and the gap closing, splitting, and merging connections are added when the corresponding entry of the assignment matrix is 1.

### SUPPLEMENTARY TEXT 2: PARAMETER OPTIMIZATION

We optimized the parameters using Ray Tune https://www.ray.io/ray-tune [4] with `BasicVariantGenerator` (random search) and `OptunaSearch` [5]. Ten rounds of 10 parallel parameter estimation were conducted for each methods, where the maximum allowed gap size was scanned from 0 to 1. The parameter ranges and initial values were set as follows.

#### Distance cutoffs and drift parameter $d$

Let $\mathcal{E}$ be the set of all given training connections with all the entry $(x, y) \in \mathcal{E}$ aligned so that $x$ exists in an earlier frame than $y$ does. The range of values and the initial value for the distance cutoff were set as $[1.5 \times q^{50\%}, 1.5 \times q^{99.9\%}]$ and $1.5 \times q^{90\%}$, respectively, where $q^{a\%}$ is the $a$ percentile of all connection distances $\{\|x - y\|_2 \,|\, (x, y) \in \mathcal{E}\}$. The cost cutoffs were then set as the square of those values. When optimizing the parameters in Eq. (4) and (5), the initial values of the distance cutoffs were set as those with the highest connection Jaccard index in the squared centroid Euclidean distance cases.

For the cell migration dataset, the range of values and the initial value for the parameter $d_j$ $(j = 1, 2)$ was set as $[m_j - s_j, m_j + s_j]$ and $m_j$, respectively, where $m_j$ and $s_j$ are the mean and the standard deviation of the values $\{(y_j - x_j) | (x, y) \in \mathcal{E}\}$.

#### Feature weight $w$

The range of values and the initial value of the parameter $w$ were set to $[0, 10]$ and 0, respectively.

#### Overlap cost function

The range of values and the initial value of the parameter $A$ were set to $[0.01, 0.5]$ and 0.01, respectively. The cost cutoff was fixed to $-\log(0.01)$. The distance cutoffs were estimated as previously, and costs are regarded as infinity for the points whose centroid distance is larger than the cutoff.

### SUPPLEMENTARY FIGURES

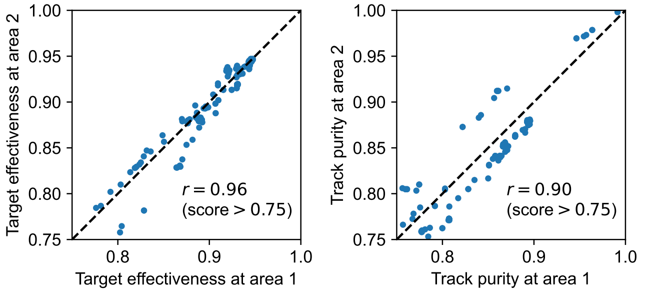


FIG. S1. The tracking scores for the mouse epidermis dataset at different regions with varied distance cutoffs [Fig. 2(b)]. The black broken lines indicate the equal scores. The Pearson correlation coefficient $r$ for data with the score values $> 0.75$ is noted in the plot.

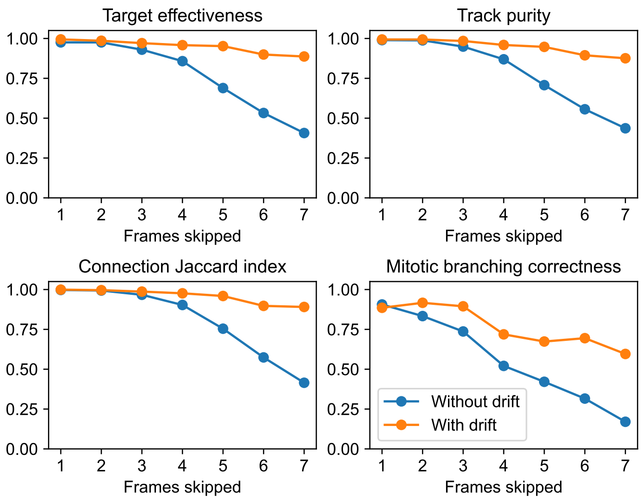


FIG. S2. All tracking scores for the cell migration dataset.

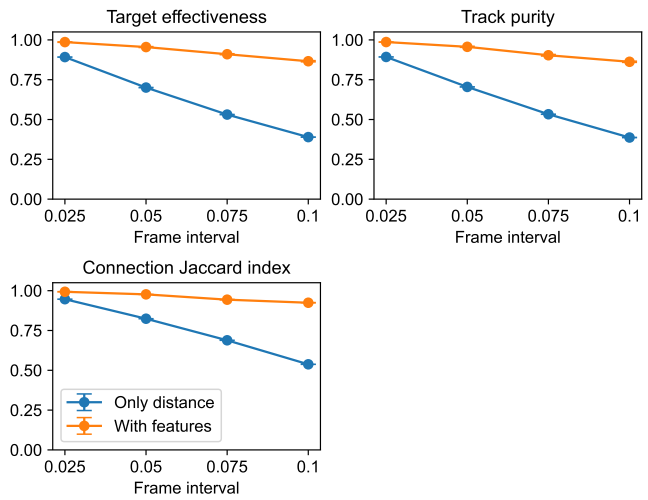


FIG. S3. All tracking scores for the colored particles dataset.
